# The role of Musashi-1 in CEP290 c.2991+1655A>G cryptic exon splicing in Leber Congenital Amaurosis

**DOI:** 10.1101/2021.08.04.454918

**Authors:** Daniele Ottaviani, Amelia Lane, Katarina Jovanovic, Jessica C. Gardner, Paul E. Sladen, Kwan L. Hau, Anna Brugulat Panes, Rosellina Guarascio, Alison J. Hardcastle, Michael E. Cheetham

## Abstract

Human photoreceptors maximise alternative exon splicing to generate a unique set of gene isoforms. Conversely, the inclusion of a cryptic exon caused by the c.2991+1655A>G deep intronic change in *CEP290* occurs in the human retina leading to Leber Congenital Amaurosis (LCA10). The RNA-binding protein Musashi-1 (MSI1) is a key component of alternative splicing in the developing mouse retina. Here we investigated the role of MSI1 in human photoreceptor-specific splicing and its potential role in *CEP290* aberrant splicing disease. Alternative splicing was studied using human induced pluripotent stem cell derived 3D retinal organoid and RPE RNA-seq datasets and several photoreceptor gene isoforms were identified. Their temporal expression was resolved in control 3D retinal organoids in comparison to development and differentiation markers. Morpholino knockdown of *MSI1* in control retinal organoids reduced the expression of several photoreceptor differentiation markers and the inclusion of photoreceptor-specific exons. Nonetheless, *MSI1* knockdown in homozygous CEP290 c.2991+1655A>G LCA10 retinal organoids did not affect the inclusion of the LCA10-associated cryptic exon. These results show that while MSI1 is important for photoreceptor alternative splicing and homeostasis, it is not a major driver of the recognition of the CEP290 cryptic splice site and the manifestation of LCA10.

**HIGHLIGHTS:**

▪ The human retina expresses a unique set of gene isoforms
▪ Musashi-1 regulates alternative splicing in 3D human retinal organoids
▪ Musashi-1 knockdown in 3D retinal organoids affects gene splicing and homeostasis in photoreceptors
▪ Musashi-1 may regulate alternative splicing of cryptic exons in retina but not in LCA10

## INTRODUCTION

The human retina is highly specialised, multi-layered sensory neural tissue with tightly regulated gene expression, especially during differentiation and development. The ability to model human inherited retinal disease through patient stem cell derived retinal organoids (RO) has created unprecedented opportunities to understand key processes occurring during retinal development and degeneration. ROs can be developed from human induced Pluripotent Stem Cells (hiPSC) obtained from patients and healthy controls thus allowing the modelling of rare and complex retinal diseases in a human context *in vitro* (Lancaster and Knoblich, 2014). In 2016, ROs were used to study the most common autosomal-recessive form of Leber Congenital Amaurosis (LCA), which is caused by the c.2991+1655A>G deep intronic change in the *CEP290* gene (CEP290-LCA or LCA10) (Parfitt et al., 2016). CEP290 is localised at both the centrosome and cilia and is regarded as an essential gatekeeper during transport of cargo across the transition zone of photoreceptor primary cilia (Craige et al., 2010). The c.2991+1655A>G change introduces a hypomorphic cryptic splice-site in *CEP290* that leads to the retention of a 128 bp pseudo-exon in the mature transcript to a premature stop codon p.Cys998* and a net loss of about 50% of the wildtype (WT) CEP290 mRNA in fibroblasts (den Hollander et al., 2006; Dulla et al., 2018; Ruan et al., 2017). Although the level of *CEP290* mRNA aberrant splicing is too low in most tissues to cause syndromic disease, the inclusion of the pseudo-exon in retina increases to over 80% of the total *CEP290* transcript in patient derived ROs, leading to a greater reduction in functional CEP290 protein potentially explaining the retinal susceptibility to LCA10 (Dulla et al., 2018; Parfitt et al., 2016). The increase in *CEP290* aberrant splicing correlated with the inclusion of retinal specific exons of *BBS8* and *RPGR*, suggesting this switch might be related to photoreceptor differentiation and specialisation (Parfitt et al., 2016). Therefore, we wanted to understand the molecular machinery that regulates splicing in the retina, timing of splice switching to retinal isoforms, and aberrant splicing such as CEP290-LCA in LCA10.

In 2016, Murphy and colleagues found that novel transcript isoforms were expressed in mouse retina and characterised the transcriptome. They identified the RNA-binding protein (RBP) Musashi-1 (MSI1) as a driver of alternative splicing (AS) and retinal-specific exons were found to be enriched in MSI1 binding motifs downstream of the splicing donor site (Murphy et al., 2016). MSI1, an RBP which targets UAG motif-enriched regions, is named after the Japanese swordsman and rōnin Miyamoto Musashi who is often portrayed holding two swords in his belt in a V-shape layout. Similarly, Drosophila knockouts for *MSI1* develop two bristles in their compound eye “ommatidia” thus resembling Musashi’s swords (Nakamura et al., 1994). This highlights a key role for *MSI1* in eye development, as also demonstrated later in mouse retina (Susaki et al., 2009). Recently, *Msi1* and its paralog *Musashi-2* were shown to be required for photoreceptor morphogenesis and survival in mice (Sundar et al., 2020). Moreover, a role for both MSI1 and MSI2 has been reported in stem cell renewal and in cancer onset, progression and metastasis (Kudinov et al., 2017; Okano et al., 2005; Wuebben et al., 2012).

Recently, Ling and colleagues performed an extensive analysis of many of the RNA-seq datasets available in literature on mouse tissues and on the human Genotype-Tissue Expression (GTEx) project with two additional datasets from the human retina (Ling et al., 2020). This revealed a high degree of AS in neuronal cells, particularly in the retina. The retina transcriptome has a distinctive splicing pattern and expresses at least 30 specific, previously unannotated, in-frame exons and micro-exons. Furthermore, MSI1 was confirmed as a driver of retinal splicing, together with PTPB1, and *MSI1* was found to be expressed mainly in rod photoreceptors thus making this mechanism likely photoreceptor specific.

Overall, these studies support the view that human photoreceptors maximise alternative exon splicing, leading to a highly specialised transcriptome during development and differentiation. This raises the possibility, however, that the retina could be particularly vulnerable to intronic mutations, compared to other tissues, as observed for the CEP290-LCA variant (Parfitt et al., 2016) and for variants within the introns of *ABCA4* that lead to Stargardt Disease (Khan et al., 2020). The mRNA processing mechanisms causing this retinal sensitivity to deep intronic variants resulting in aberrant splicing are still unknown.

In this study, we exploited RNA-seq from ROs and Retinal Pigment Epithelium (RPE) to identify the RO specific transcriptome. We aimed to correlate the inclusion of photoreceptor-specific exons and the CEP290-LCA pseudoexon with that of other AS exons, and to investigate the role of MSI1. We developed ROs to define the timing of the AS program during human retinal development *in vitro*. We then exploited Morpholino-mediated *MSI1* gene silencing. We show MSI1 is indispensable for the execution of the photoreceptor-specific splicing program in human retinal tissue but has a limited impact on the inclusion of the CEP290-LCA cryptic exon.

## RESULTS

### Identification of alternative splicing events unique to retinal organoids

We recently developed an RNA-seq transcriptome for 3D ROs from a healthy donor (Lane et al., 2020); RPE from the same hiPSC line were differentiated in parallel and also sequenced by RNA-seq. The rMATS pipeline compares two experimental conditions and identifies patterns of exon skipping, alternative 5’ and 3’ splice site usage, mutually exclusive exons and retained introns (Shen et al., 2014). It was used to explore and compare the RO and RPE datasets to identify differentially expressed exons (AS events) and the potential photoreceptor transcriptome. We found 11,606 exons were differently spliced in ROs and many that were not previously annotated are reported here (Supplementary Table 1). Among the identified exons, 5074 were significantly upregulated in ROs while, 6532 were downregulated (Supplementary Table 1). Despite being enriched in photoreceptors, ROs contain many cell types of the human retina; namely Muller-Glia, retinal ganglion cells, interneurons (bipolar, horizontal, amacrine) and RPE (Collin et al., 2019). Furthermore, exons could be shared with other tissues as well. Therefore, we compared the AS exons with those from mice (Murphy et al., 2016) in order to narrow down the analysis to the common photoreceptor AS events and to test for exon conservation. The exon coordinates shortlisted by Murphy *et al*. were aligned by synteny to the human genome thus allowing filtering of our dataset for AS events that matched. We found 102 exons enriched in the human retina have mouse orthologues (Supplementary Table 2). From the top 20 most significant upregulated and downregulated orthologues displaying the highest inclusion difference between RO and RPE (Figure 1A, B), we initially focused on a previously unannotated micro-exon of 17bp in length in the *IMPDH1* gene (*IMPDH1* exon-14), shown in Figure 1A. In 2016, IMPDH1 exon-14 expression in the human retina was reported by VastDB, an extensive database of alternative spicing events in vertebrates (Tapial et al., 2017). Among the 20 shortlisted exons, *IMPDH1* exon-14 was validated by Murphy and colleagues by RT-PCR in mouse retina, together with *CC2D2A* (Murphy et al., 2016). Although awaiting validation in humans, *IMPDH1* exon-14 showed the strongest MSI1-dependent expression in mice (Murphy et al., 2016). Therefore, we selected the *IMPDH1*-exon 14 as a reporter to test when the AS switch occurs in RO development and to investigate if MSI1 is part of this mechanism of exon selection in the human retina.

**Figure 1.**
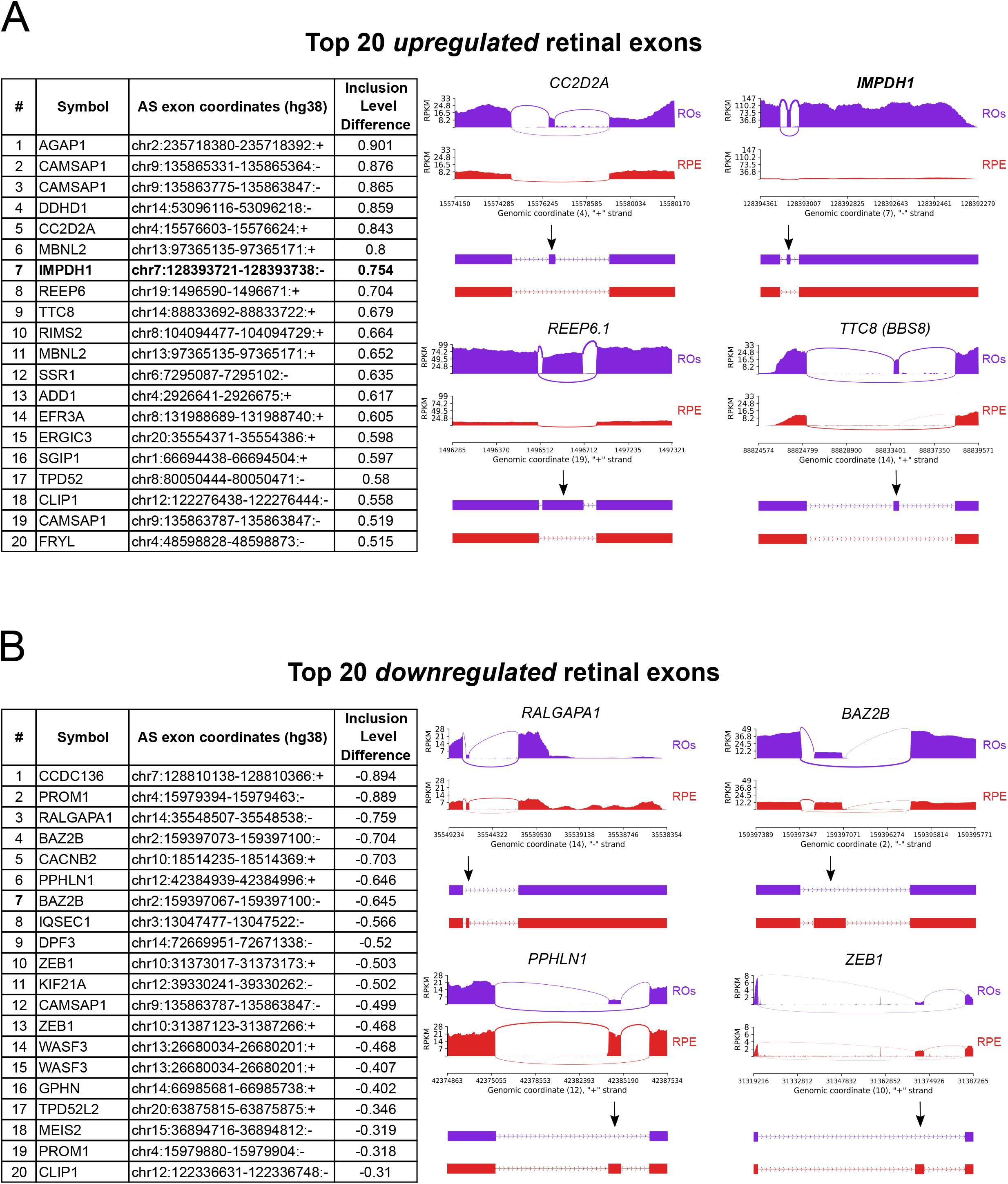
Alternatively spliced genes in 3D retinal organoids (ROs) compared to the RPE. **(A)** Top 20 upregulated AS genes in ROs compared to the RPE. The *IMPDH1* exon 14 is highlighted in bold and was selected as a reporter for AS events. Right panel illustrates exon inclusion in ROs compared to RPE (*CC2D2A*, *IMPDH1*, *REEP6.1* and *TTC8*) **(B)** Top 20 downregulated AS genes in ROs compared to the RPE.Right panel illustrates exons more common in RPE compared to ROs (*RALGAPA1*, *BAZ2B*, *PPHLN1* and *ZEB1*).

### Retinal-specific isoforms appear during retinal organoid development

The expression of known markers of retinal differentiation was investigated in control ROs at different time-points of differentiation *in vitro*; day (D) 20, 40, 50, 60, 70, 90, 100, 120, 150 and 180 (Figure 2). RT-PCR showed that specifiers of retinal progenitor cells, *PAX6* and *VSX2/Chx10*, are concordantly and constantly expressed during RO differentiation and development from about D20 onwards (Figure 2A). In contrast, the photoreceptor-specific transcription factors *CRX*, *NRL* and *NR2E3* were detected at later timepoints showing an increase from D50, D70 and D90 onwards, respectively (Figure 2A). We next performed quantitative real-time PCR (qPCR) over the time-course of RO differentiation and extended our analysis to day 200. There was exponential increase in expression for the photoreceptor associated genes when the data were plotted on a semi-Log transformed scale (Figure 2B). The results showed a similar trend of exponential increase for the expression of *CRX*, *NRL* and *NR2E3* starting from D20 in ROs (Figure 2B), except for *PAX6* and *VSX2*, which were relatively constant.

**Figure 2.**
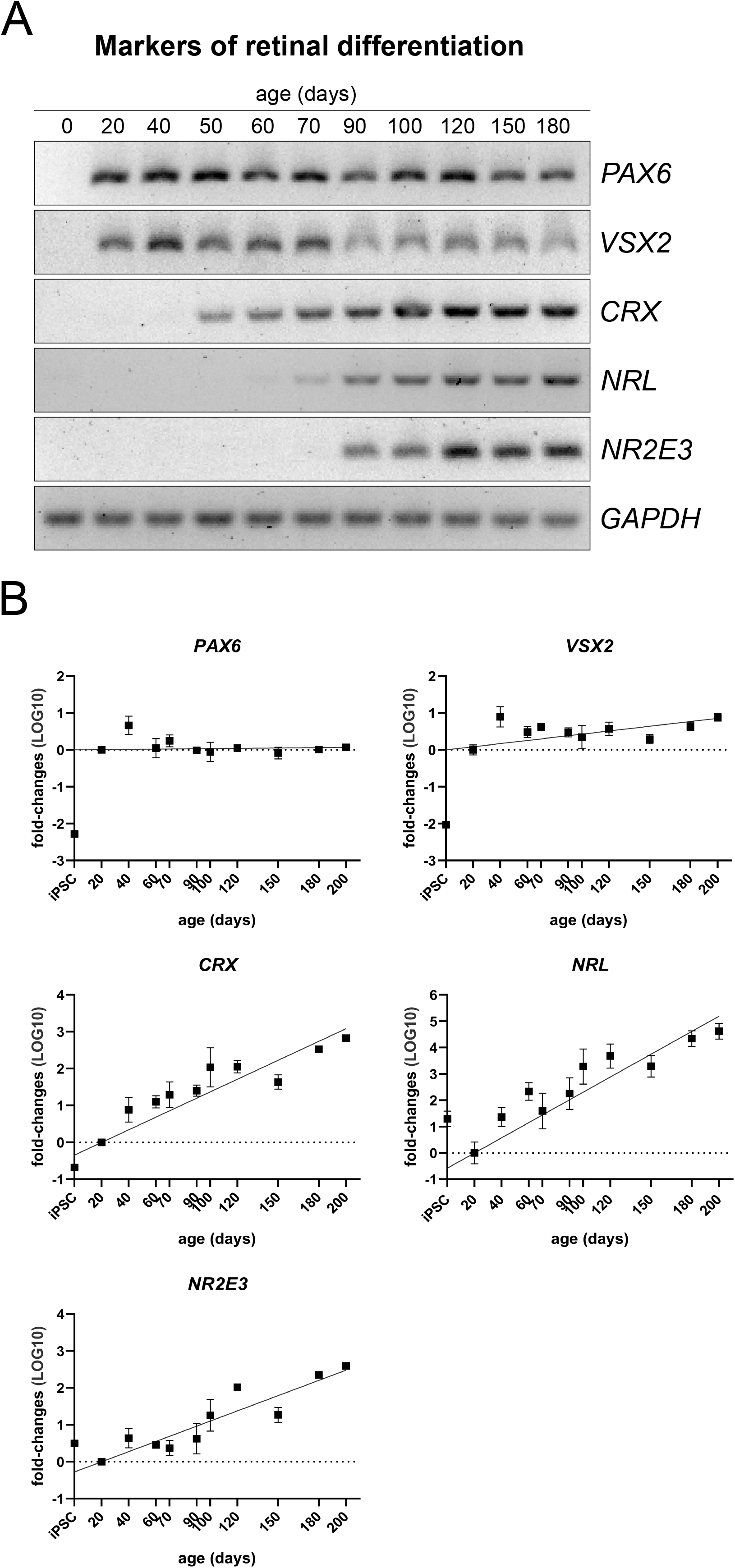
Temporal resolution of retina-specific exons in retinal organoids. **(A)** RT-PCR of control RO retinal differentiation markers *PAX6*, *VSX2*, *CRX*, *NRL* and *NR2E3*. **(B)** qPCR semi-log plots showing their temporal expression in control retinal organoids at day 20, 40, 70, 90, 100, 150, 180, 200 (mean of n≥3 experiments ± SD for each time point).

These stages of RO development were used to characterize when retinal-specific exons are alternatively spliced in human retina. Retinal-specific isoforms have been previously described for *BBS8/TTC8*, *REEP6* and *RPGR* and mutations in these retinal exons have been linked to retinal degeneration (Arno et al., 2016; Kirschner et al., 1999; Murphy et al., 2015). Therefore, the expression of retinal-specific isoforms *BBS8* exon-2A, *REEP6.1*, *RPGRorf15* were investigated with the addition of *IMPDH1* exon-14 (Figure 3). There was consistent increase of *RPGRorf15* transcripts from D40, which preceded that of *REEP6.1* and *BBS8* exon-2A, which were first detected at D50 (Figure 3A). Transcripts containing Exon 14 of *IMPDH1* were detectable at D20 at the first stages of RO differentiation, suggesting the retinal splicing program might be active early during retinal development *in vitro* (Figure 3A). We performed a more resolved qPCR analysis and found *BBS8* exon-2A, *REEP6.1*, *RPGRorf15* and *IMPDH1* exon-14 are all detectable at D20 (Figure 3B). Notably, there was a close fit with markers of photoreceptor differentiation, *CRX*, *NRL* and *NR2E3*, whose transcription rate rise exponentially from D20. These results suggest the retinal splicing program is initiated early during RO development *in vitro* and qPCR yields the appropriate resolution of the inclusion of retinal-specific isoforms that seem to closely follow photoreceptor differentiation and development in ROs.

**Figure 3.**
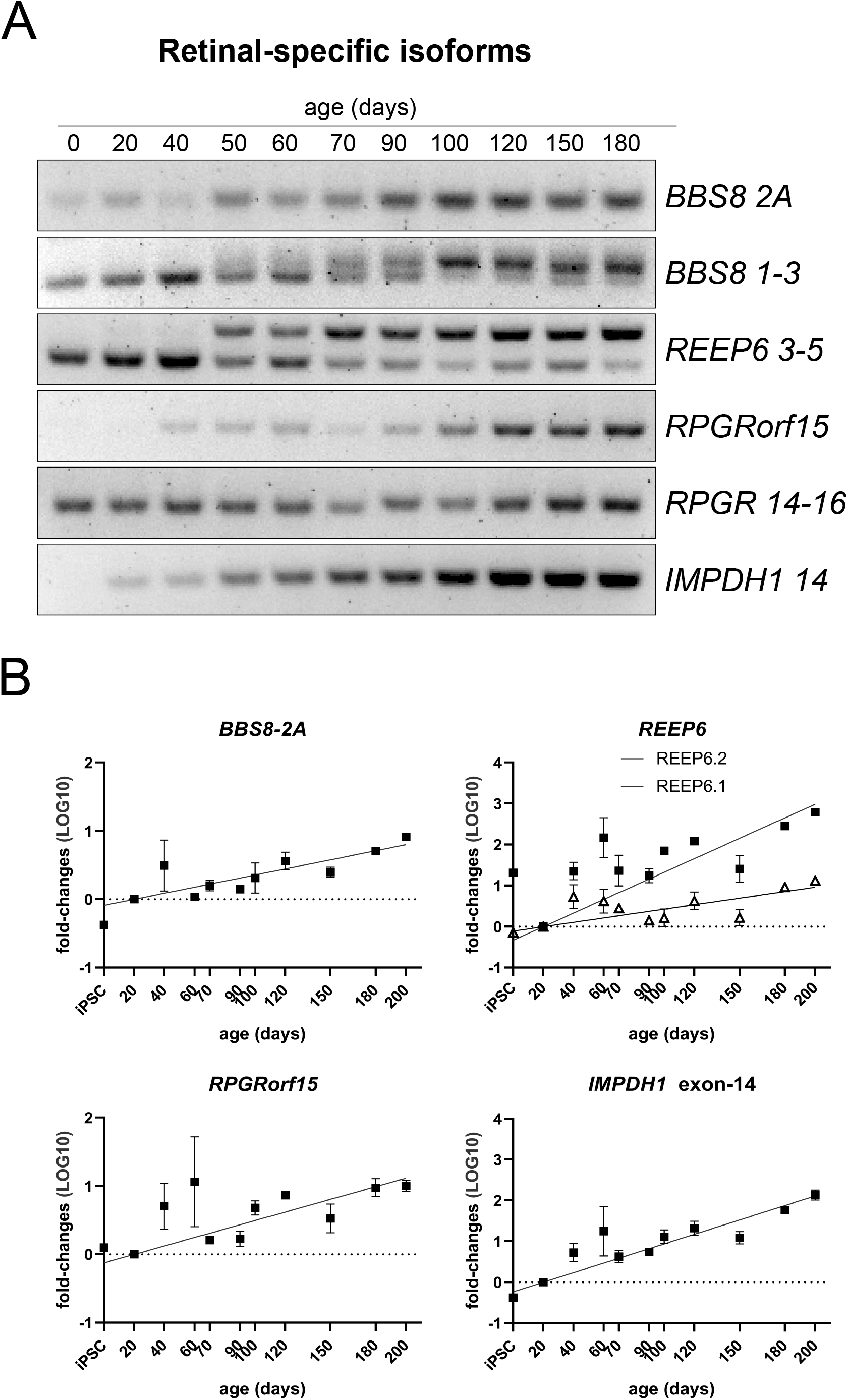
Temporal resolution of retina-specific exons in retinal organoids. **(A)** RT-PCR of control RO retinal-specific exons in *BBS8* (*BBS8* exon-2A), *REEP6.1* (upper band), *REEP6.2* (lower band), *RPGRorf15* and *IMPDH1* exon-14. **(B)** qPCR semi-log plots showing their temporal expression in control ROs at day 20, 40, 70, 90, 100, 150, 180, 200 (mean of n≥3 experiments ± SD for each time point).

### MSI1 knockdown affects photoreceptor gene expression

We used an antisense oligonucleotide (Morpholino, *MSI1*-MO) to knockdown the expression of *MSI1* in mature ROs to study the role of MSI1 in retinal homeostasis. Control ROs were treated twice with 10 μM of *MSI1*-MO or 10 μM of a control non-specific MO over a 5-day period at D195 of development. This window of treatment was selected as the photoreceptors have completed development and were likely to yield a high qPCR signal-to-noise ratio after treatment. *MSI1* downregulation was analysed by quantifying the mRNA level by qPCR (Figure 4A) and by plotting fold-changes in gene expression. This confirmed a statistically significant 50% reduction in the level of *MSI1* transcript (Figure 4A). The global markers of neuronal cell fate *VSX2* (Figure 4B) and *PAX6* (Figure 4C) were not altered by *MSI1* knock-down. In contrast, genes related to the photoreceptor cell lineage were significantly decreased, with the rod photoreceptor-specific genes *NRL* (Figure 4D) and *NR2E3* (Figure 4E) showing a 60% reduction and more affected than the cone-rod marker *CRX*, which showed a 40% decrease (Figure 4F). This suggests reduction of *MSI1* could affect photoreceptor homeostasis and survival, with possibly a greater impact on rod function when compared to cones.

**Figure 4.**
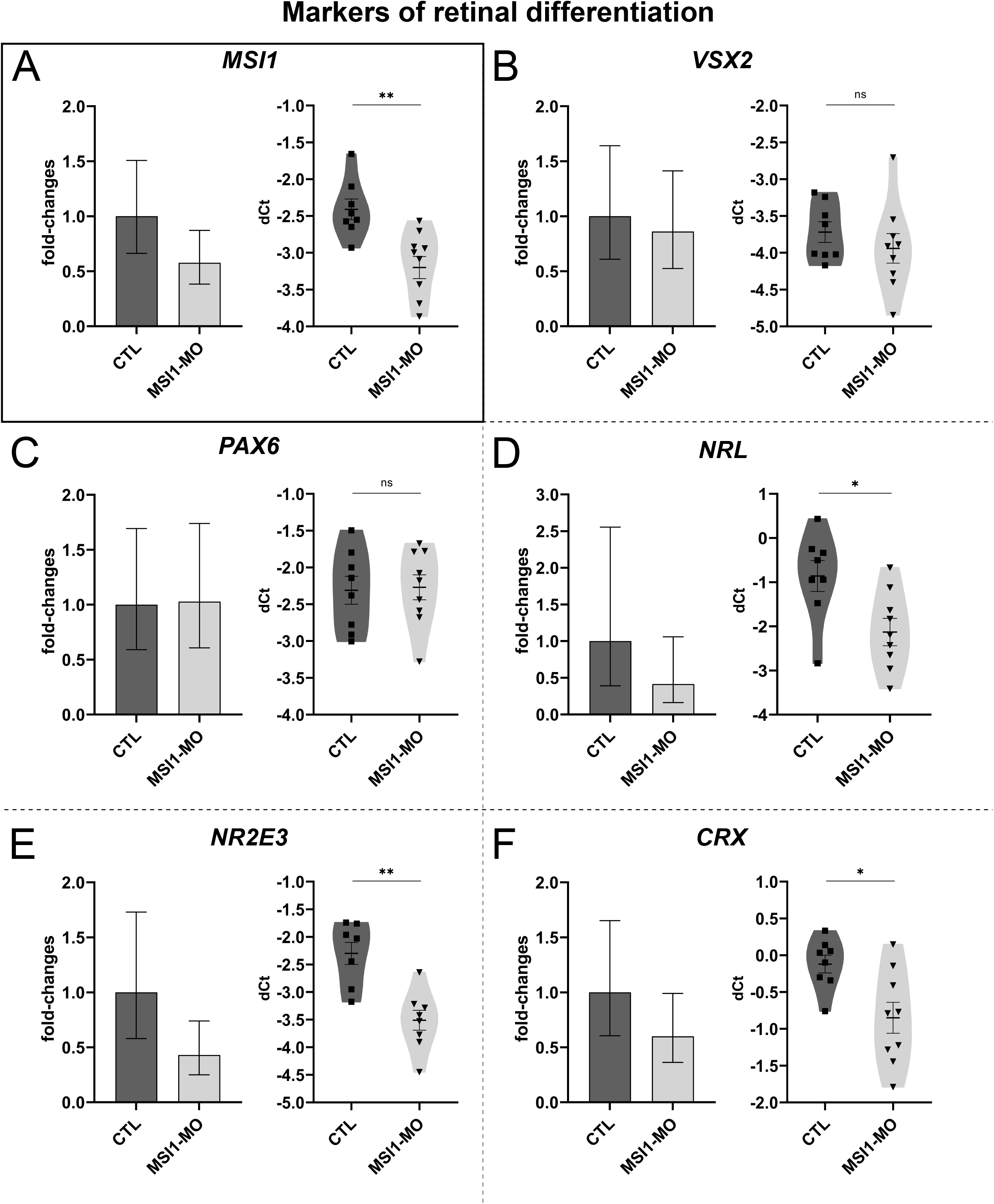
qPCR analysis for markers of retinal differentiation in control organoids treated with the MSI1-Morpholino. qPCR analysis of markers of retinal differentiation *VSX2, PAX6*, *NRL, NR2E3* and *CRX* (Panel A-F) in control ROs treated with 10 μM of the MSI1-Morpholino (*MSI1*-MO) for 5 days. The MSI1 knockdown effect is highlighted with solid square as reference for working *MSI1*-MO. For each gene, the graph on the right shows fold-changes compared to the mean of control-MO treated (CTL) ROs; individual replicates are plotted as reciprocal of the dCt values used to calculate the fold-changes. Statistics was performed on dCt values by Student t-test with the GraphPad Prism 5 software. (mean of n=9 organoids ± SD. Each point represents an individual organoid. *P≤0.05, **P≤0.01,***P≤0.001, unpaired, two-tailed, Student’s t-test).

### MSI1 is important for alternative splicing in human retinal organoids

We investigated the effect of *MSI1*-MO treatment on retina-specific exons that have previously been reported as containing consensus sites for MSI1 (Ling et al., 2020), starting with *IMPDH1* exon-14. The inclusion of *IMPDH1* exon-14 was analysed and was found to be significantly reduced by approximately 50% (Figure 5A). Furthermore, the retinal-specific exon 30 in the *CC2D2A* gene was also reduced, but exon 24 of *CC2D2A*, another retinal-specific exon in the same transcript was not reduced (Figure 5B). *CASK* and *ADGRV1* retinal exons were significantly decreased by 60% and 90%, respectively, in the *MSI1* knockdown ROs (Figure 5C, D), whereas *DOC2B* was not affected by *MSI1* knockdown as shown previously in mice (Figure 5E) (Ling et al., 2020). Surprisingly, we observed that *REEP6.1* splicing was also affected by MSI1 reduction (Figure 5F), which has not been previously reported as containing consensus sites for MSI1 (Ling et al., 2020). However, not all of the exons with MSI1 consensus sites were affected by the *MSI1*-MO in this assay. While *IMPDH1* exon-14, *CC2D2A* exon-30, *ADGRV1* and *CASK* show a splicing pattern strongly dependent on *MSI1* expression, an alternative mechanism likely exists for other retinal exons such as *CC2D2A* exon-24 and *DOC2B*.

**Figure 5.**
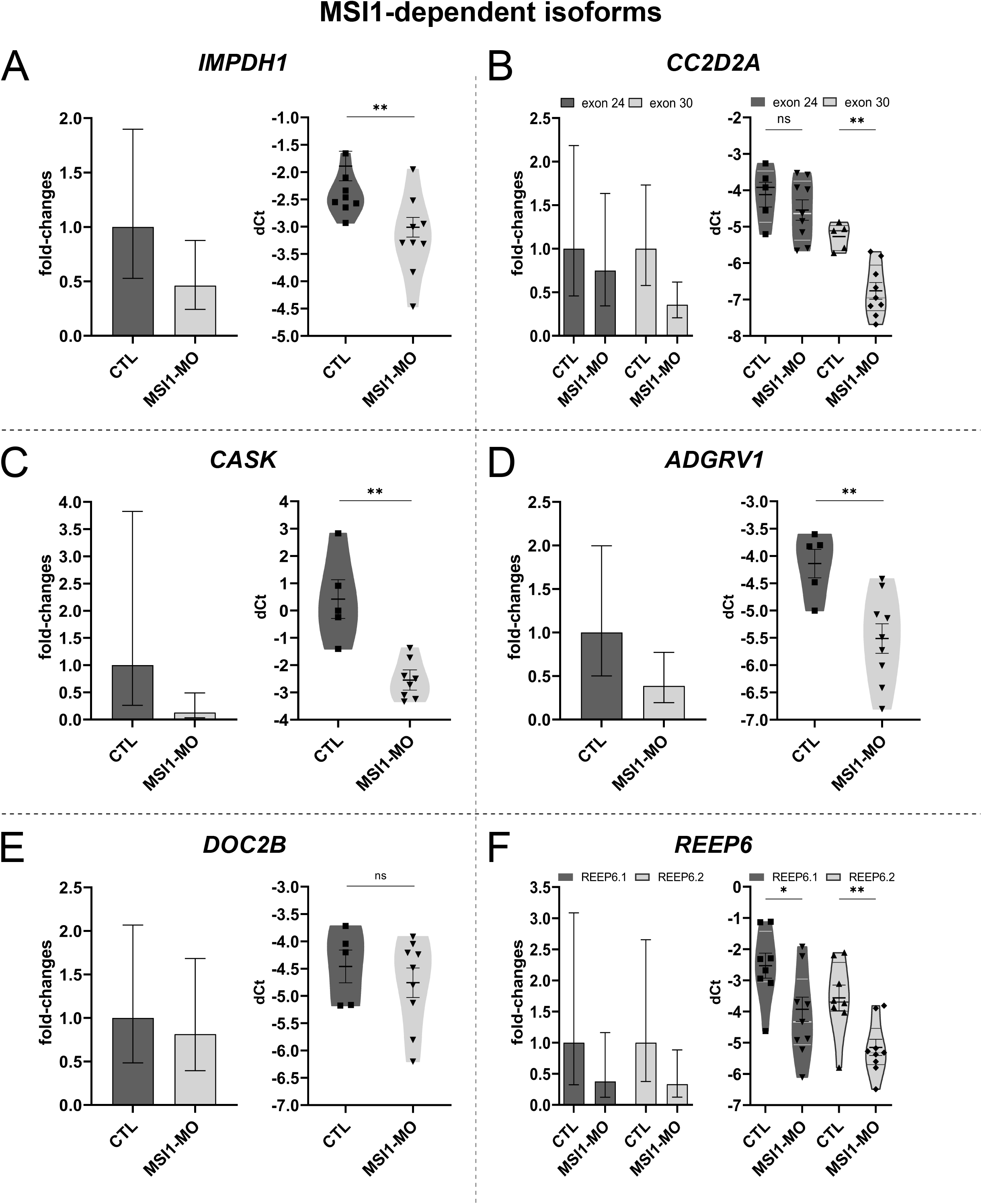
qPCR analysis for retinal-specific exons of control organoids treated with the MSI1-Morpholino. qPCR analysis of retinal-specific exons in *IMPDH1* exon-14, *CC2D2A* exon 24 and 30, *CASK*, *ADGRV1*, *DOC2B, REEP6.1* and *REEP6.2* (Panel A-F) in control ROs treated with the MSI1-Morpholino treated with 10μM of MSI1-MO for 5 days (*MSI1*-MO). For each gene, the graph on the right shows fold-changes compared to the mean of control-MO treated (CTL) ROs; individual replicates are plotted as reciprocate of the dCt values used to calculate the fold-changes. Statistics was performed on dCt values by Student t-test with the GraphPad Prism 5 software. (mean of n=9 organoids ± SD. Each point represents an individual organoid. *P≤0.05, **P≤0.01,***P≤0.001, unpaired, two-tailed, Student’s t-test).

### MSI1 drives photoreceptor-specific splicing in human retinal organoids

Murphy and colleagues identified the MSI1-driven AS in mouse retina; whereas, Ling and collaborators suggested this AS program is rod specific and built an extensive database of AS in human tissue, named ASCOT (Ling et al., 2020). Our AS analysis matches ASCOT findings for the most relevant retinal pathways and disease-associated genes (Figure 6A and Supplementary table 3).

**Figure 6.**
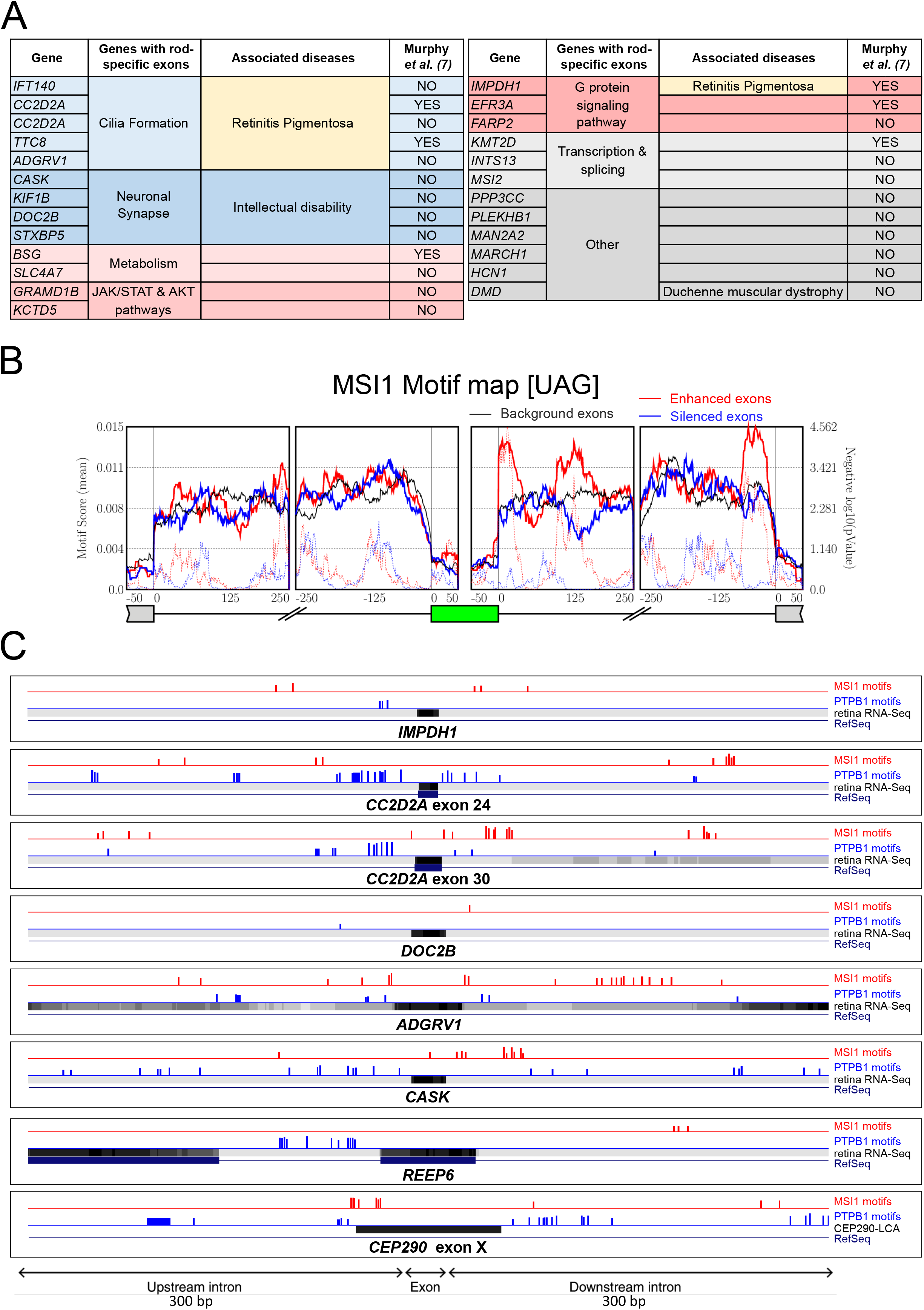
Analysis of selected retinal-specific exons and CEP290-LCA cryptic exon. **(A)** AS events in selected genes in ROs that match those identified by Ling and colleagues in ASCOT (Ling et al., 2020). **(B)** rMAPS2 analysis of the enrichment in MSI1 binding motifs in upregulated (Enhanced, red line) and downregulated, skipped (Silenced, blue line) and up/downstream of the 3’ and 5’ splice sites, respectively. **(C)** RBPmaps analysis displayed on the UCSC Genome Browser shows the locations of MSI1 binding motifs (red peaks) and PTPB1 binding motifs (blue peaks). The peak heights reflect different significance of the RBPmaps results (the higher the peak, the smaller the p-value, p-value cutoff < 0.01). The RefSeq is shown along with the retina RNA-Seq used to build ASCOT (Ling et al., 2020), as some of the exons are not yet annotated. IMPDH1 exon-14, CC2D2A exon 24 and 30,DOC2B, ADGRV1, CASK, REEP6.1 and CEP290-LCA cryptic exon are reported.

Furthermore, we investigated the MSI1-dependence of the AS events we identified in ROs by using rMAPS2. rMAPS2 takes the rMATS AS results and looks for enrichment in RBP binding motifs among up- and down- regulated isoforms. These analyses revealed that many exons upregulated in ROs are enriched in tandem MSI1-binding motifs in the 50 bp and 125 bp regions downstream to the 5’ splice site (Figure 6B). This is in agreement with a defined MSI1-enriched region of 200 bp downstream of the AS exons (14). Interestingly, there were also MSI1 motif peaks at the 3’ splice site of downstream exons which could be AS.

To investigate further the discordant behaviour of the AS exons we investigated in the previous section, we compared locations of their MSI1 binding motifs by using RBPmaps (Paz et al., 2014) (Figure 6C). RBPmaps identifies binding motifs for RBPs in sequences of interest. As the novel retinal transcriptome shows conservation across mammals (Ling et al., 2020; Tapial et al., 2017), we hypothesised that regulatory regions would be under the same evolutionary constraints and applied the RBPmaps conservation filter, which considers the degree of sequence conservation between human and mouse. *IMPDH1* exon-14, *CC2D2A* exon 30, *ADGRV1* and *CASK* meet the requirements of the majority of MSI1-driven exons, specifically being enriched in MSI1 binding motifs in close proximity to the donor splice site and in PTBP1 binding motifs upstream to the acceptor splice site (Ling et al., 2020; Murphy et al., 2016). Conversely, *CC2D2A* exon 24 has MSI1 tandem motifs further downstream to the 5’ splice site while *DOC2B* displays a single motif. Our data suggest these sites might be inactive with respect to regulation of these retinal exons. Surprisingly, *REEP6.1* has putative binding motifs for MSI1; however, they are at the suggested limit of the MSI1 binding range (Figure 6C) (Ling et al., 2020). Moreover, the *REEP6.2* isoform was also reduced, such that this could be independent of retinal specific splicing. This shows a more complex mechanism might regulate splicing in retina, possibly in different cell types, or that another exon which is common to both *REEP6.1* and *REEP6.2* might be affected.

Intriguingly, we noticed the cryptic exon X CEP290-LCA displays long-range downstream MSI1 binding sites similar to those of *REEP6.1* (Figure 6C). Therefore, it is unlikely they could regulate the CEP290-LCA AS splicing pattern. Nonetheless, CEP290-LCA displays a repeated MSI1 motifs cluster upstream of the 3’ splice site that is conserved, despite the CEP290-LCA pseudoexon not being a natural exon. This is in accordance with the ASCOT analyses which found that high levels of MSI1 expression can activate non-conserved, otherwise non-expressed, cryptic exons such as in the *RDX* and *GDAP1* genes. Ling et al. found these cryptic exons are unusually enriched in MSI1 sites upstream to the 3’ splice site, as well as downstream to the 5’ splice site (Ling et al., 2020). The retina has a high level of *MSI1* gene expression and the CEP290-LCA pseudoexon seems to closely fit the MSI1 motifs cluster for cryptic exons at the 3’ splice site. This prompted us to investigate if MSI1 could affect the retinal mis-splicing in CEP290-LCA.

### The role of MSI1 in CEP290 c.2991+1655A>G cryptic exon splicing in LCA10

The CEP290-LCA10 cryptic exon displays long range MSI1 binding motifs downstream of the cryptic splice site and putative binding motifs for both MSI1 and PTPB1 upstream (Figure 6B). This led us to consider MSI1 as a potential regulator of CEP290-LCA10 aberrant splicing, and the pseudo-exon as a novel target. To test this hypothesis, we differentiated homozygous c.2991+1655A>G CEP290-LCA ROs and treated them with *MSI1*-MO to test if MSI1 plays a role in the aberrant splicing of *CEP290*. We performed a 5-day treatment at D195 of CEP290-LCA RO development, and in control ROs, and quantified the amount of *MSI1* and *IMPDH1* exon-14, as a readout of a successful MSI1 functional knockdown (Figure 7). The *MSI1* transcript was decreased following *MSI1*-MO treatment in CEP290-LCA ROs, and the inclusion of *IMPDH1* exon 14 was reduced similar to control ROs, which confirmed effective reduction of MSI1-dependent splicing (Figure 7A). In contrast, there was only a small non-significant reduction in the inclusion of the CEP290-LCA10 cryptic exon. Furthermore, this was not accompanied by an increase in the WT (exon 26-27) *CEP290* transcript (Figure 7A, B), as was observed when using splice switching therapeutic antisense oligonucleotides for LCA10 (Dulla et al., 2018; Parfitt et al., 2016). Therefore, it appears that MSI1 is not a major driver of the photoreceptor specific switch in CEP290-LCA cryptic exon inclusion and that it is likely other factors have a role.

**Figure 7.**
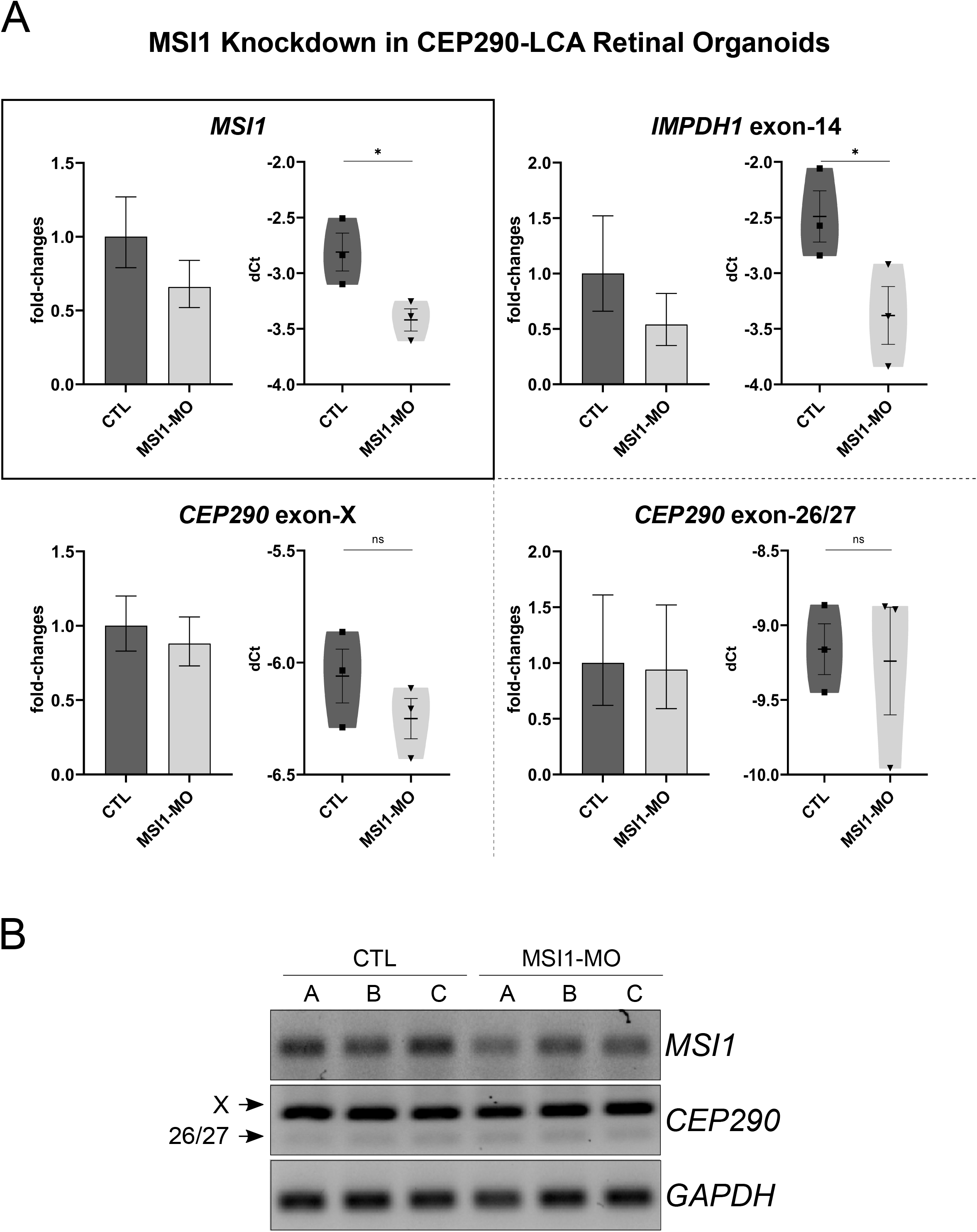
qPCR analysis of CEP290-LCA retinal organoids treated with the MSI1-Morpholino. **(A)** qPCR analysis of *MSI1*, *IMPDH1* exon 14, CEP290 cryptic and canonical isoforms in D195 CEP290-LCA retinal organoids treated with 10μM of MSI1-MO for 5 days. The MSI1 knockdown effect is highlighted with a red square and used as a reference for successful knockdown with the *MSI1*-MO. For each gene, the graph on the right shows fold-changes compared to the mean of control-MO treated (CTL) ROs; individual replicates are plotted as reciprocate of the dCt values used to calculate the fold-changes. Statistics was performed on dCt values by Student t-test with the GraphPad Prism 5 software. (mean of n=3 organoids ± SD. Each point represents an individual organoid. *P≤0.05, **P≤0.01,***P≤0.001, unpaired, two-tailed, Student’s t-test). **(B)** RT-PCR image analysis of *MSI1*, *IMPDH1* exon 14, CEP290 cryptic and canonical isoforms in CEP290-LCA retinal organoids treated with MSI1-MO.

## DISCUSSION

The impact of alternative splicing on fundamental biological processes, as well as health and disease has become increasingly evident thereby driving the characterization and investigation of the underlying mechanisms. Sequencing techniques have become progressively affordable and have allowed unprecedented insights into AS. The most advanced RNA-seq techniques can capture the dynamic nature of the mammalian transcriptome, which can facilitate gene evolution under pressure. For this reason, the relevance of the plethora of AS identified has been debated, whether it is mostly a mechanism of regulation to determine which gene isoforms are switched off by nonsense mediated decay, or if it actively produces functional alternative transcriptome (Blencowe, 2017; Tress et al., 2017). The development of bioinformatic pipelines and cross-comparisons between published datasets have made it possible to navigate this evolutionary “noise” to extract AS events that are conserved in mammals and of interest to human pathophysiology. Among the AS databases, VastDB (Tapial et al., 2017) and, more recently, ASCOT (Ling et al., 2020), have been key to our understanding of AS in the brain and retina. In particular, the brain shows the most abundant degree of AS and has been under the spotlight since the pioneering studies on microexons and their implication in autism (Irimia et al., 2014). The retina, as part of the CNS, has recently come into focus and an increasing body of evidence points to genetic variants affecting AS associated with inherited blinding conditions; namely, *ABCA4*, *CEP290*, *DYNC2H1* (Khan et al., 2020; Parfitt et al., 2016; Vig et al., 2020).

The c.2991+1655A>G deep intronic change in *CEP290* predominantly affects the retina via an unknown AS mechanism, which exacerbates CEP290 aberrant splicing in photoreceptors (Dulla et al., 2018; Parfitt et al., 2016). Unfortunately, *in vivo* humanised animal models of the c.2991+1655A>G LCA10 variant are of limited use, as they do not fully recapitulate *CEP290* aberrant splicing and the human LCA10 phenotype (Garanto et al., 2013). Therefore, we generated hiPSC-derived ROs from a CEP290-LCA patient and healthy control as models of retinal development *in vitro*. We defined the retinal transcriptome by comparing ROs and RPE. Then, we exploited published datasets to cross-compare AS events of interest in ROs and found they are in accordance with the AS Databases. We observed the retinal AS program starts as early as day 20 of ROs development and increases exponentially up to 200 days in culture *in vitro*. We also found MSI1 is an essential driver of AS for many genes in the human retina and defined the constrains for its activity.

Musashi-1 has been shown to be a master regulator of photoreceptor AS both in mouse and in humans (Ling et al., 2020; Murphy et al., 2016; Sundar et al., 2020); therefore, we investigated the dependence of shortlisted exons on MSI1. Knockdown of *MSI1* in ROs shows a direct effect on those exons that strictly met the previously defined MSI1 dependencies (Ling et al., 2020; Murphy et al., 2016). However, *MSI1* knockdown had effects that extended beyond AS alone, through the reduction of *NRL*, *NR2E3* and *CRX*, that has implications for retinal development, photoreceptor function and survival. At present, it is not clear whether the mechanism for this reduction is through a direct effect on these transcripts or by acting via other factors that influence their expression, but it highlights the importance of MSI1 for retinal homeostasis and potential uninvestigated roles. This is in partial agreement with a mouse double conditional knockout for *Msi1* and *Msi2*, which shows defects in photoreceptor morphogenesis, due to impaired outer segment development (Sundar et al., 2020). The conditional pan-retinal *Msi1* and *Msi2* knockouts developed a laminated retina suggesting no major defects in stemness or in photoreceptor specification; however, there were alterations in retinal cell survival, including defects in the neuroblastic layer as early as P5. We showed instead that key photoreceptor transcription factors were affected by the *MSI1* knockdown in human retinal organoids, with reduction of *CRX*, *NRL* and *NR2E3* and this could influence retinal development or maintenance. Interestingly, the mouse *Msi1* conditional KO alone had a less pronounced effect on AS or retinal function and survival, perhaps indicating a greater importance for MSI1 in human retina.

Moreover, we found MSI1 seems dispensable for *DOC2B*, *CC2D2A* exon-24 AS and has little effect on the cryptic exon splicing in CEP90-LCA. This suggests that an alternative scenario may exists in human retina and that other RBPs might be involved. For example, PRPF31 has been shown to influence retinal AS in ROs and RPE (Buskin et al., 2018), and several splicing factors have been identified as retinal disease genes. More recently, a study on AS during organ development in mammals highlighted the RBP QKI being at the hinge of AS in heart and brain development (Chen et al., 2021; Mazin et al., 2021). Indeed, the rMAPS2 analysis we performed showed a significant peak for QKI just upstream to the 3’ splice sites (Supplementary Figure S1). However, this might be because many AS events we identified share a broader neuronal expression, rather than an exclusively retinal one, such that the link with CEP90-LCA is still missing. Nonetheless, iPSC-derived ROs are a valuable model to faithfully recapitulate conditions affecting AS in human retina, with special relevance to intronic variants and for developing potential therapies (Cideciyan et al., 2019, 2021; Khan et al., 2020; Parfitt et al., 2016).

## EXPERIMENTAL PROCEDURES

### Bioinformatic analysis

RNA-Seq analysis of ROs and RPE was carried out as described previously (Lane et al., 2020). Three ROs and RPE replicates from the control BJ line (ATCC CRL-2522) were harvested at D150. The RNA was extracted with the RNeasy micro kit (QIAGEN) following the manufacturer’s instructions, followed by paired-end sequencing at 100 million read depth for each sample (Illumina, Otogenetics, Atlanta, GA, USA). Raw.fastq sequences were cleaned from any residual sequencing adapter using cutadapt with parameters -m 20 and -e 0.1. Fragments were then aligned to the human genome (build 38, Ensembl version 92) using STAR. BAM files were then used by rMATS to detect differentially spliced exons in retinal organoids (ROs) compared to the RPE data with settings –novelSS to enable detection of novel splice sites (unannotated splice sites). Sashimi plots were made using the function rmats2sashimiplot.

### Online resources and tools

We exploited different online resources and tools. Namely, VastDB https://vastdb.crg.eu/wiki/Main_Page, ASCOT http://ascot.cs.jhu.edu/, rMAPS2 http://rmaps.cecsresearch.org/ and RBPmap http://rbpmap.technion.ac.il/ and the UCSC Genome Broswer https://genome.ucsc.edu/.

### Differentiation of iPSCs to RPE and ROs

RPE was differentiated as in (Schwarz et al., 2015), while ROs were differentiated according to the protocol described by Zhong and collaborators (Zhong et al., 2014); with modifications (de Bruijn et al., 2020; Lane et al., 2020). In brief, iPSCs were grown to 95% confluence in E8 medium. Colonies were gently scraped in gentle dissociation buffer (ThermoFisher Scientific) to form embryoid bodies (EBs). EBs were transitioned to neural induction medium in the presence of blebbistatin (Sigma) before seeding at a density of approximately 20 EBs per cm^2^. Emerging transparent pouches of neuroepithelium were isolated using a needle or scalpel and cultured in Maturation Medium (3:1 v/v of DMEM: F12, 2% B27 supplement, 1% Non-Essential Amino Acid, 1% Penicillin-Streptomycin, 10% Fetal Bovine Serum (Labtech), 100 μM Taurine, 2 mM GlutaMAX) until day 70, then changed to Retinal Maturation Medium 2 (3:1 v/v of DMEM: F12, 1% N2 supplement, 1% Non-Essential Amino Acid, 1% Penicillin-Streptomycin, 10% Fetal Bovine Serum (Labtech), 100 μM Taurine, 2 mM GlutaMAX). Media was supplemented with 1 μM retinoic acid from day 50 to day 70, then changed to 0.5 μM.

### MSI1 knockdown, RNA Extraction and qPCR

The Standard Control and *MSI1* Morpholino (Supplementary Table S4) were designed and synthesized by Gene Tools (Gene Tools, LLC). ROs were mock- or MSI1 treated with 10 μM the *MSI1* Morpholino for 5 days with an additional boost at day 3, following the manufacture protocol.

RNA was extracted from cell lines and retinal organoids using the RNeasy Micro Kit (Qiagen, Hilden, Germany) following the manufacturer’s instructions. Then, 50 ng of RNA were converted to cDNA by the Tetro cDNA Synthesis Kit (Bioline, London, UK) and a mixture of oligo-dT and random hexamer primers.

qPCR analysis was performed using the LabTaq Green Hi Rox (Labtech) following manufacturer’s instructions on a QuantStudio 6 Flex Real-Time PCR System (Applied Biosystems) and primers listed in Supplementary Table S4.

Relative gene expression levels were determined with the ΔΔCt method. Statistics and plots were made with GraphPad Prism v.8 (GraphPad Software, Inc.)

## Data and Code Availability

The RO and RPE datasets are available at GEO GSE148300 and GSE180531, respectively.

## ACKNOWLEDGEMENTS

This work was supported by funding from Retina UK, the Wellcome Trust, Fight for Sight, the NC3Rs and MRC.

## AUTHOR CONTRIBUTIONS

D.O., A.L., K.J., J.C.G., P.E.S., K.L.H., A.B.P. and R.G. performed the experiments and/or analysed the data. D.O., A.L., K.J., A.J.H. and M.E.C. conceived the hypothesis and designed the experiments. D.O., A.J.H. and M.E.C. drafted the manuscript. All authors edited the draft manuscript.

## DECLARATION OF INTERESTS

The authors declare no competing interests.

## SUPPLEMENTAL INFORMATION

**Supplementary Table S1. rMATS Alternative splicing results.** Exons with significant difference in the inclusion levels in retinal organoids (ROs) compared to the RPE are highlighted.

**Supplementary Table S2. rMATS Alternative splicing results and filtered for orthologues**. Exons with significant difference in the inclusion levels in retinal organoids (ROs) compared to the RPE shortlisted for events identified in mice (Murphy et al., 2015).

**Supplementary Table S3. Alternative splicing analysis**. Exons with significant difference in the inclusion levels in retinal organoids (ROs) compared to the RPE and matched to those shortlisted in ASCOT (Ling et al., 2020).

**Supplementary Table S4.** Morpholino and Primer sequences for RT and qPCR.

**Supplementary Figure S1.** rMAPS2 analysis of the enrichment in QKI binding motifs in upregulated (Enhanced, red line) and downregulated, skipped (Silenced, blue line) and up/downstream of the 3’ and 5’ splice sites, respectively

## Notes

### Competing Interest Statement

The authors have declared no competing interest.

