## Supplementary figures and images for "The role of Musashi-1 in CEP290 c.2991+1655A>G cryptic exon splicing in Leber Congenital Amaurosis"

### Supplemental Figure S1

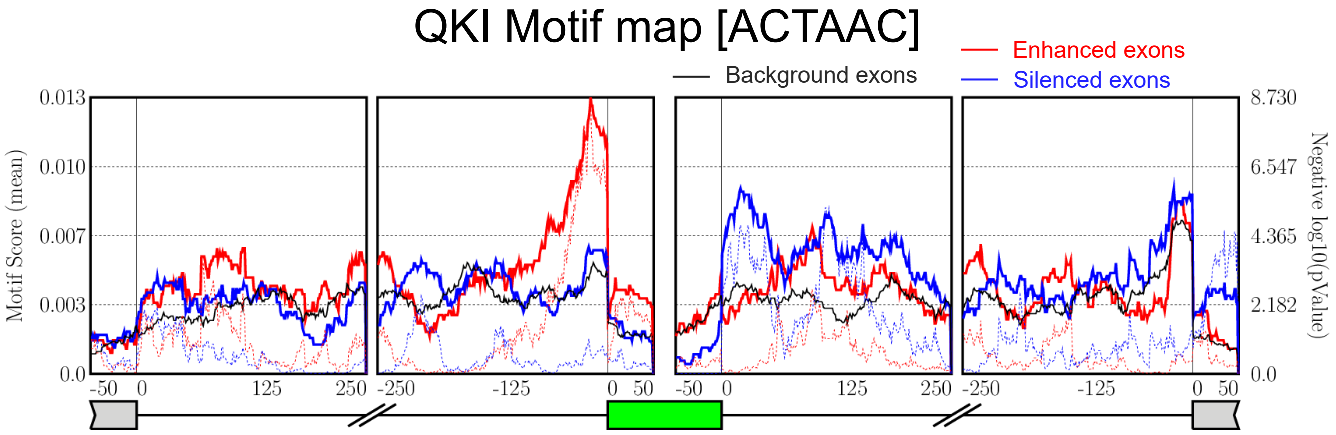
