## Supplemental Table S4 for "The role of Musashi-1 in CEP290 c.2991+1655A>G cryptic exon splicing in Leber Congenital Amaurosis"

### Table S4 Morpholino and Primer sequences for RT and qPCR.

| Target | Forward |
|---|---|
| **Morpholino** | |
| ***MSI1* targeting transcription initiation (ATG)** | CCTCTTACCTCAGTTACAATTTATA |
| **Gene Tools Standard Control** | CGTCAGTCTCCATCGGGAGCC |

| **Primers for RT-PCR only** | | |
|---|---|---|
| **Target** | **Forward** | **Reverse** |
| ***BBS8 TOTAL*** | GGAGCTATTTTAGGCGCAGG | TTTCAAAGACGTTCCAGGGC |
| ***REEP6 TOTAL*** | TCCTGTCCTGGTTCCCTTTC | GGCTGCTTCACTTGTCCTTC |
| **qPCR primer sequences** | | |
| **Target** | **Forward** | **Reverse** |
| ***MSI1*** | GACTCAGTTGGCAGACTACGC | ACCCCTGGATCTCTTGGTCAG |
| ***IMPDH1 (retinal isoform)*** | CATGGCCTGCACTCCTATACC | GAGCTGGAGAACCCGTAGTG |
| ***RPGRorf15*** | GGATTTTCATGACGCAGCCAG | CTACCTCTTGCTCCTCTATTCCA |
| ***BBS8 (retinal isoform)*** | CATCAGGCAGCTTGGATCTTAA | TTCAAAGACGTTCCAGGGCG |
| ***REEP6.1 (retinal isoform)*** | CAGGAACGTCTTGCAGGTCC | CTTGAGGTCAGCTTCCAGGG |
| ***REEP6.2*** | CTGTTCCTAAGGCACCACGG | GCTTGGCTTGACGTTCCTGG |
| ***CRX*** | TTTGCCAAGACCCAGTACC | GTTCTTGAACCAAACCTGAACC |
| ***NRL*** | CACTGACCACATCCTCTCGG | GAGGGTTCCCGCTTTACCTC |
| ***NR2E3*** | TCTTCAAGCCAGAGACGCG | CTCAAAGACGGGAGGAGCAG |
| ***VSX2 (Chx10)*** | GTGGCTACTGGGGATGCAC | TCCTGCTCCATCTTGTCGAG |
| ***PAX6*** | AACGATAACATACCAAGCGTGTC | GTCTGCCCGTTCAACATCCT |
| ***CC2D2A (retinal isoform 1)*** | CCAATATCAGTTCTGAAGGCTCA | GGTTTTGGGCTGTCACCTTC |
| ***CC2D2A (retinal isoform 2)*** | CCATTCGAGAAAAGATGAGTAACA | GGAAACTTTAAGGCACATTCAGC |
| ***DOC2B (retinal isoform)*** | GCCAAGGTGCCAAAATTGATG | GAGGGTCTCGTTCCATGTGG |
| ***ADGVR1 (retinal isoform)*** | AGCAGCCATCAACATTACCA | GAGTCTTTTCTAGTCTGGGAGG |
| ***CASK (retinal isoform)*** | CAATACTGGACGTGGAGCCTC | GACGCTGAAGAGATTCATAGGC |
| ***GAPDH*** | CCCCACCACACTGAATCTCC | GGTACTTTATTGATGGTACATGACAAG |
| ***ACTB*** | CCAACCGCGAGAAGATGA | CCAGAGGCGTACAGGGATAG |
